# Marginal Returns and Levels of Research Grant Support among Scientists Supported by the National Institutes of Health

**DOI:** 10.1101/142554

**Authors:** Michael Lauer, Deepshikha Roychowdhury, Katie Patel, Rachael Walsh, Katrina Pearson

## Abstract

The current era of worsening hypercompetition in biomedical research has drawn attention to the possibility of decreasing marginal returns from research funding. Recent work has described decreasing marginal returns as a function of annual dollars granted to individual scientists. However, different fields of research incur varying cost structures. Therefore, we developed a Grant Support Index (GSI) that focuses on grant activity code, as opposed to field of study or cost. In a cohort of over 71,000 unique scientists funded by NIH between 1996 and 2014 we analyzed the association of grant support (as measured by annual GSI) with 3 bibliometric outcomes, maximum Relative Citation Ratio (which arguably reflects a scientist’s most influential work), median Relative Citation Ratio, and annual weighted Relative Citation Ratio (which is more dependent on publication counts). We found that for all 3 measures marginal returns decline as annual GSI increases. Thus, we confirm prior findings of decreasing marginal returns with higher levels of research funding support.

## Introduction

In a recent essay on the “systemic flaws” besetting biomedical research, Alberts et al called on funding agencies to “be sensitive to the total numbers of dollars granted to individual laboratories, recognizing—although different research activities have different costs — at some point, returns per dollar diminish.”^1^ A growing body of literature supports the claim that “at some point, returns per dollar diminish,” or more precisely, that as funding levels or personnel support^2^ for any given laboratory increases, *marginal* (that is incremental) returns decline.^3^ However, as Alberts et al recognized, it may be problematic to focus solely on dollars precisely because different types of research entail varying cost structures. We therefore developed a “Grant Support Index” (GSI) that focuses on grant activity code, as opposed to field of study or cost, and using this measure attempted to <u>confirm other reports</u> of decreasing marginal returns with increased research funding.^3-6 7,8^

## Methods

### Study sample and description of grant support

We identified NIH Research Project Grants (“RPGs”) that use currently available grant activity codes and were funded between 1996 and 2014 approximately 2,500 grantee organizations, including domestic and private institutions of higher education, research institutions and hospitals. The data were primarily extracted from the Information for Management, Planning, Analysis, and Coordination (IMPAC II) database for grants and Research and Development contracts, and public interface is known as eRA Commons. Over the period analyzed, 71,936 unique scientists served as Principal Investigators (PIs) or Multiple Principal Investigators (MPIs) on NIH RPG grants. For each grant-PI/MPI– Fiscal Year combination, we assigned GSI points using a scheme that NIH staff derived without knowledge of research outcomes. A trans-NIH Extramural Activities Working Group (EAWG) subcommittee, under the leadership of Jon Lorsch (Director, NIGMS), held several dedicated meetings that involved dozens of NIH program and grants management staff to develop the GSI (originally designed the “<u>Research Commitment Index</u>” or RCI). We began with assigning a standard R01 grant 7 points, and then, based on that benchmark, assigned more or fewer points to other mechanisms. The agreed-upon point scheme is shown in Table 1; only after the scheme was agreed to did we begin analyzing outcomes.

**Table 1:** Grant Support Index (GSI) point values for NIH Research Project Grants. These were values assigned in the summer of 2016 and reported out in earlier in 2017.

| Activity Code | Single PI point assignment | Multiple PI point assignment |
| --- | --- | --- |
| UM1, UM2 | 11 | 10 |
| **Subprojects** under multi-component awards | 6 | 6 |
| R01, R33, R35, R37, R56, RC4, RF1, RL1, P01, P42, UC4, UF1, UH3, U01, U19, DP1, DP2, DP3, DP4 | 7 | 6 |
| R00, R21, R34, R55, RC1, RC2, RL2, RL9, UG3, UH2, U34, DP5 | 5 | 4 |
| R03, R24, P30, UC7 | 4 | 3 |

We gathered data on Howard Hughes Medical Institution (HHMI) funding support from the HHMI web-site and manually matched HHMI awardees to NIH awardees by name and institution.

### Outcomes

The “Relative Citation Ratio” (RCR) is a bibliometric measure developed by the NIH Office of Portfolio Analysis that compares citation influence of articles while accounting for field of study and time of publication.^9^ Briefly, RCR compares actual and expected citation rates by assembling and inspecting a “co-citation network,” papers cited alongside the paper of interest. The RCR has been shown to correlate well with expert opinion.^9^ We used the NIH Scientific Publication Information Retrieval and Evaluation Systems^10^ (“SPIRES”) to link NIH PI’s to publications; we obtained RCR values for each publication from the NIH Office of Portfolio Analysis.

We limited our analyses to papers published during the study period of 1996-2014 (N=1,027,462). We focused on 3 outcomes:

- Maximum RCR: the highest RCR for NIH RPG-supported paper linked to a scientist’s RPG grants; in communications with the authors, some have endorsed the importance of this measure as a reflection of scientist’s most influential work. Other recent efforts have estimated individual scientists’ impacts by considering their most highly cited papers;^11^
- Median RCR; and
- Annual weighted RCR: we summed the RCR values for all papers linked to a scientist’s grants. This measure is most sensitive to publication counts, which vary by discipline.^12^In cases where more than one scientist was linked to a paper (e.g. papers that acknowledge support from multiple grants, from multi-PI grants, and from multicomponent grants), we divided and distributed the RCR values among all involved PI’s to avoid double-counting papers.

### Analyses

Our approach mirrors those of prior reports^3,4^ that specifically addressed the question of how research impact scales with funding. To examine the association between grant support and research output, we examined the association of annual GSI with the 3 RCR outcomes. We focused on annual GSI as the independent variable because all NIH budgeting and grant decisions are made on a year-to-year basis. As annual GSI and RCR values followed skewed distributions, we used natural log-transformed values throughout.

With only one independent variable, GSI, we used a Cobb-Douglas function^3^ with the form *ln(Q) ~ ln(α)* + *β(ln(K)),* where Q is output as assessed by maximum RCR, median RCR, or annual weighted RCR, *K* is input (in this case annual GSI), and β is a slope coefficient reflecting marginal returns that links the natural logarithm of input (annual GSI) with the natural logarithm of output. A value of β > 1 corresponds to increasing returns (e.g. a 10% increase in input is associated with > 10% increase in output); a value of β = 1 corresponds to constant returns (e.g. a 10% increase in input corresponds to a 10% increase in output); a value of b between 0 and 1 corresponds to decreasing returns (e.g. a 10% increase in input corresponds to < 10% increase in output); and a value of β < 0 corresponds to negative returns (e.g. as input increases, output decreases). We allowed for non-linear associations between *Q* and annual GSI by testing natural cubic spline terms with 3, 4, or 5 knots. For all models tested, all terms (linear and non-linear spline) were highly significant (with all *P* < 2 * 10^-16^). We used Akaike Information Criterions (AIC) to compare models.

Because of community concerns regarding the use of natural log-log transformations, we also derived loess regression plots of a random subsample of 20,000 scientists to show associations with annual GSI of maximum RCR and annual RCR.

All analyses were performed in R (https://www.r-project.org/).

## Results

### Sample

We identified 71,936 unique scientists who served as Principal Investigators (PI’s) on NIH Research Project Grants (including sub-projects) between 1996 and 2014. We assigned “Grant Support Index” points according to the scheme shown in Table 1; points varied by mechanism and by whether there were more than one PI for grants in which multi-PI applications were allowed. A GSI value of 7 corresponds to 1 single-PI R01 while a GSI of 21 corresponds to 3 single-PI ROls.

Table 2 shows characteristics and bibliometric outcomes of PI’s according to four categories of Grant Support Index (GSI) that correspond to ≤ 1 R01 equivalent, > 1 to ≤ 2 R01 equivalents, > 2 to ≤ 3 R01 equivalents, and > 3 R01 equivalents. Among all PI’s, 46,365 (64%) had an annual GSI of 7 or less, while 21,182 (29%) had an annual GSI between 7 and 14. Only 6% of PI’s had an annual GSI of 14 or higher.

Figures S1 – S6 show box plots of the distributions of maximum RCR, median RCR, and annual weighted RCR. These box plots demonstrate the skewed distributions.

### Grant Support and Productivity

Figure 1A shows the association of maximum RCR with annual GSI (with, as previously noted, both axes natural log-transformed to reflect the skewed distributions and the underlying Cobb-Douglas function). The rug plots along the axis demonstrate the log-linear distributions of annual GSI and maximum RCR. Increasing levels of annual GSI were associated with increasing levels of maximum RCR, but the *rate* of increase decreases with increasing annual GSI. Figure 2A illustrates this point by showing the instantaneous slope (first derivative, or instantaneous value of β) across values of annual GSI; this curve effectively shows marginal returns. Marginal returns fall as annual GSI increases; once GSI exceeds 21, marginal returns decrease to the point that any increase in input was associated with a proportionally lower increase in output.

**Table 2:** Characteristics and outputs of 71,936 NIH-supported scientists according to their annual number of RPG Grant Support Index points.

| | GSI $\leq 7$ | | | GSI $> 7$ and $\leq 14$ | | | GSI $> 14$ and $\leq 21$ | | | GSI $> 21$ | | |
| --- | --- | --- | --- | --- | --- | --- | --- | --- | --- | --- | --- | --- |
|  | *N* = 46365 |  |  | *N* = 21182 |  |  | *N* = 3450 |  |  | *N* = 939 |  |  |
| Annual Grant Support Index | 5.0 | 6.6 | 7.0 | 8.3 | 9.5 | 11.2 | 15.0 | 16.3 | 18.0 | 22.5 | 24.3 | 27.7 |
| Annual BRDPI-Adjusted \$M | 0.16 | 0.28 | 0.39 | 0.41 | 0.53 | 0.70 | 0.82 | 1.02 | 1.40 | 1.37 | 1.80 | 2.41 |
| Years of RPG Funding | 2 | 3 | 5 | 7 | 10 | 15 | 11 | 16 | 19 | 15 | 19 | 19 |
| Annual Publications | 0.29 | 0.93 | 1.77 | 1.06 | 1.70 | 2.57 | 1.88 | 2.65 | 3.80 | 2.40 | 3.48 | 4.94 |
| Annual Weighted RCR | 0.19 | 0.93 | 2.41 | 1.28 | 2.45 | 4.46 | 3.01 | 4.98 | 8.30 | 4.31 | 7.26 | 11.73 |
| Mean RCR | 0.4 | 1.0 | 1.7 | 1.1 | 1.5 | 2.1 | 1.4 | 1.9 | 2.6 | 1.6 | 2.1 | 2.7 |
| Median RCR | 0.32 | 0.79 | 1.29 | 0.75 | 1.05 | 1.43 | 0.95 | 1.21 | 1.54 | 1.02 | 1.28 | 1.60 |
| Maximum RCR | 0.61 | 2.13 | 4.97 | 3.32 | 6.26 | 12.20 | 7.91 | 13.66 | 24.62 | 12.25 | 20.40 | 34.86 |

*a b c* represent the lower quartile *a*, the median *b*, and the upper quartile *c* for continuous variables.
GSI = Grant Support Index (see Table 1)
BRDPI = Biomedical Research and Development Price Index
\$M = millions of dollars of total costs
RPG = NIH Research Project Grants
RCR = Relative Citation Ratio

**Figure 1A:**
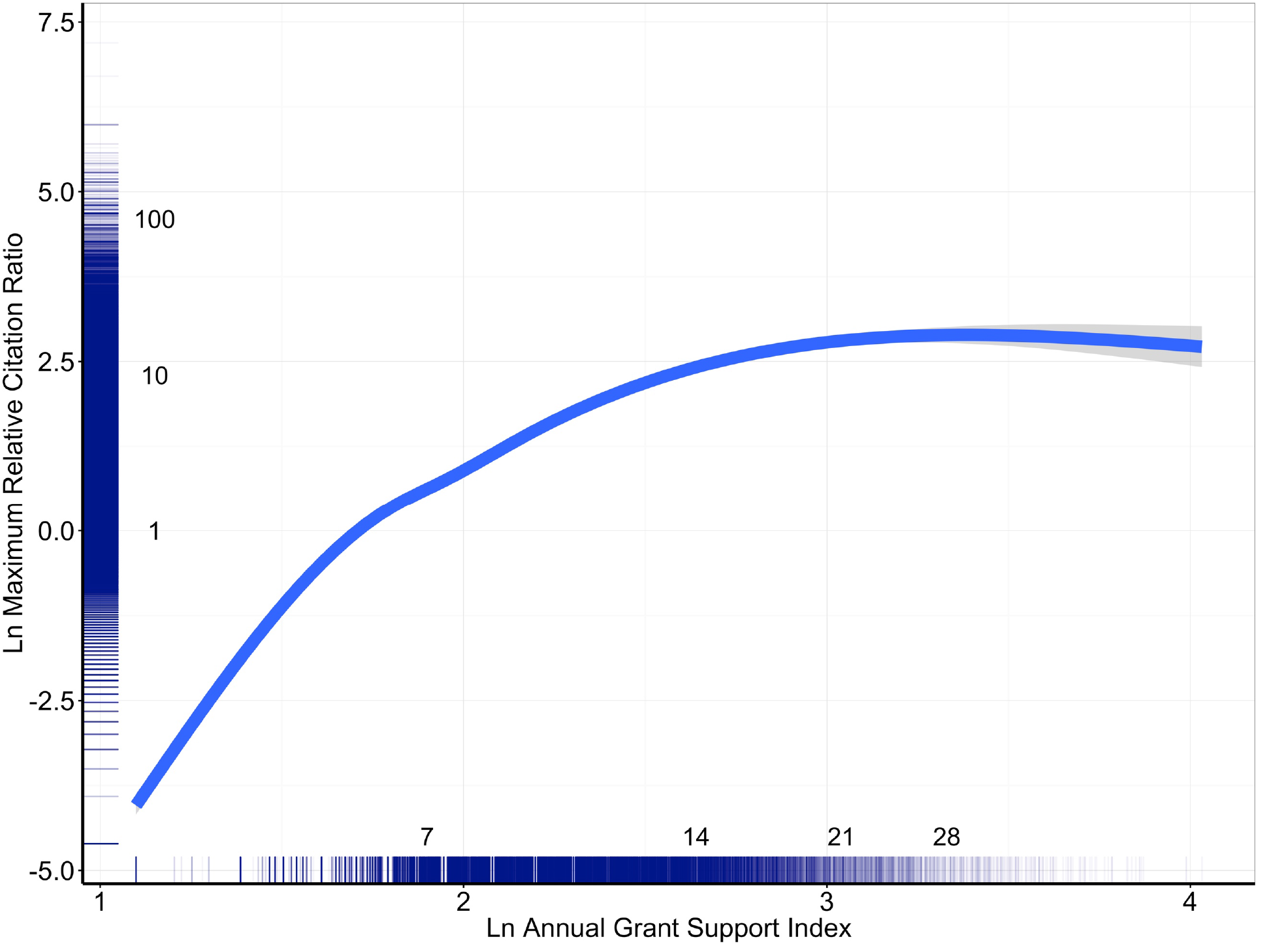
Maximum Relative Citation Ratios by level of Grant Support. Association, as assessed by spline regression, of the maximum Relative Citation Ratio (RCR) among all papers linked to a scientists’ grants with annual Grant Support Index (GSI). Due to skewed distributions, both RCR and GSI values are natural log-transformed; the numbers inside the axes represent the raw, non-transformed values. An annual GSI value of 7 corresponds to ~ 1 R01 grant, while annual GSI values of 14 and 21 correspond to 2 and 3 R01 grants. The rugplots by the axes illustrate the log-normal distributions. Note the decreasing slope of the regression curve as annual GSI increases.

**Figure 2A:**
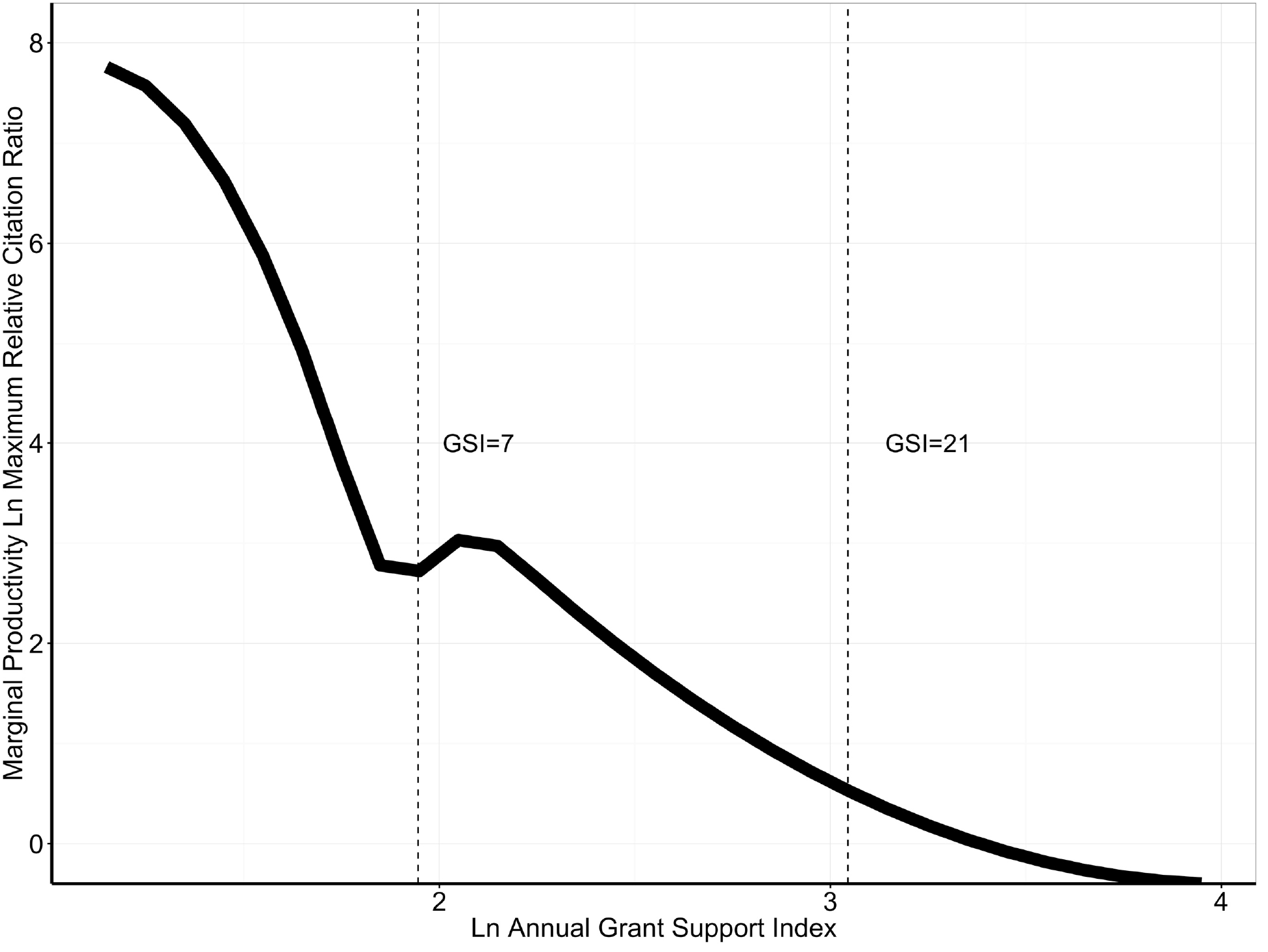
Marginal returns of maximum RCR by level of Grant Support. Association of the marginal returns of maximum Relative Citation Ratio (RCR) among all papers linked to a scientists’ grants with annual Grant Support Index (GSI). The curve shows the slope of the curve shown in Figure 1A and corresponds to the first derivative *d(ln(Q))/d(ln(K))* of the Cobb-Douglas function *ln(Q) ~ ln(α)* + *β(ln(K))*, where *Q* is output as assessed by maximum RCR and p is a coefficient linking the natural logarithm of input (K or *annual GSI)* with the natural logarithm of output, here maximum RCR. For convenience, vertical dashed lines are shown corresponding to GSI values of 7 (equivalent to 1 single-PI R01) and 21 (equivalent to 3 single-PI R01s). Marginal productivity falls as annual Grant Support Index increases.

We tested a non-transformed model using spline regression and found substantially worse fit (by AIC) than with a natural log-log model. Figure 1B is based on loess regression of a randomly chosen subsample of 20,000 scientists. Figure 2B shows that the instantaneous slope (or first derivative) of the loess regression curve becomes shallower with increasing annual GSI, reflecting a decline in marginal returns.

**Figure 1B:**
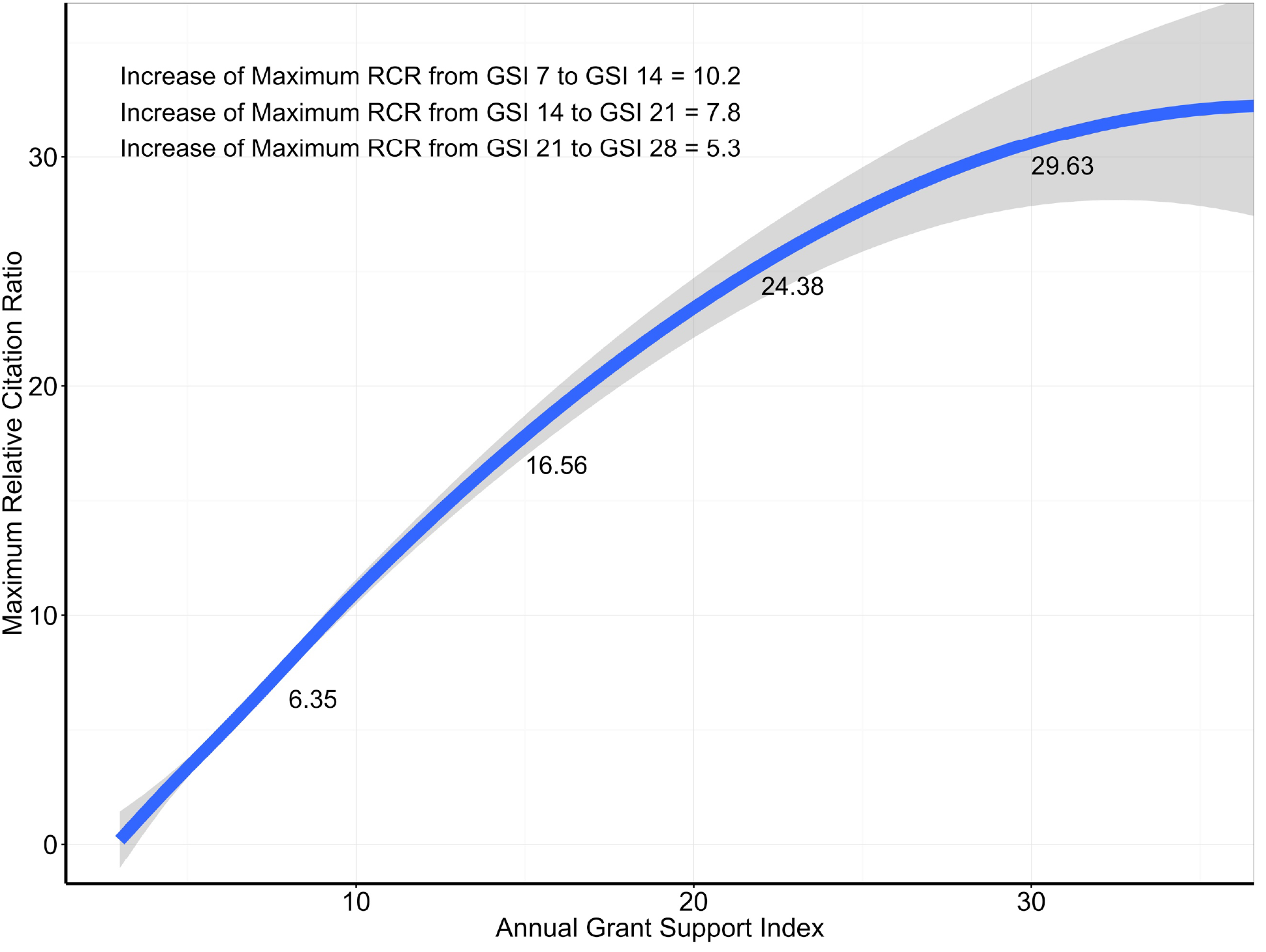
Maximum Relative Citation Ratios and Grant Support – loess regression of random subsample. Association, as assessed by loess regression in a random subsample of 20,000, of the maximum Relative Citation Ratio (RCR) among all papers linked to a scientists’ grants with annual Grant Support Index (GSI). Here values are not log-transformed. Loess fit values and changes are shown along the curve and in the upper left-hand corner to illustrate decreasing incremental returns with increasing levels of grant support. The decreasing instantaneous slope (by first derivative) is shown in Figure 2B.

**Figure 2B:**
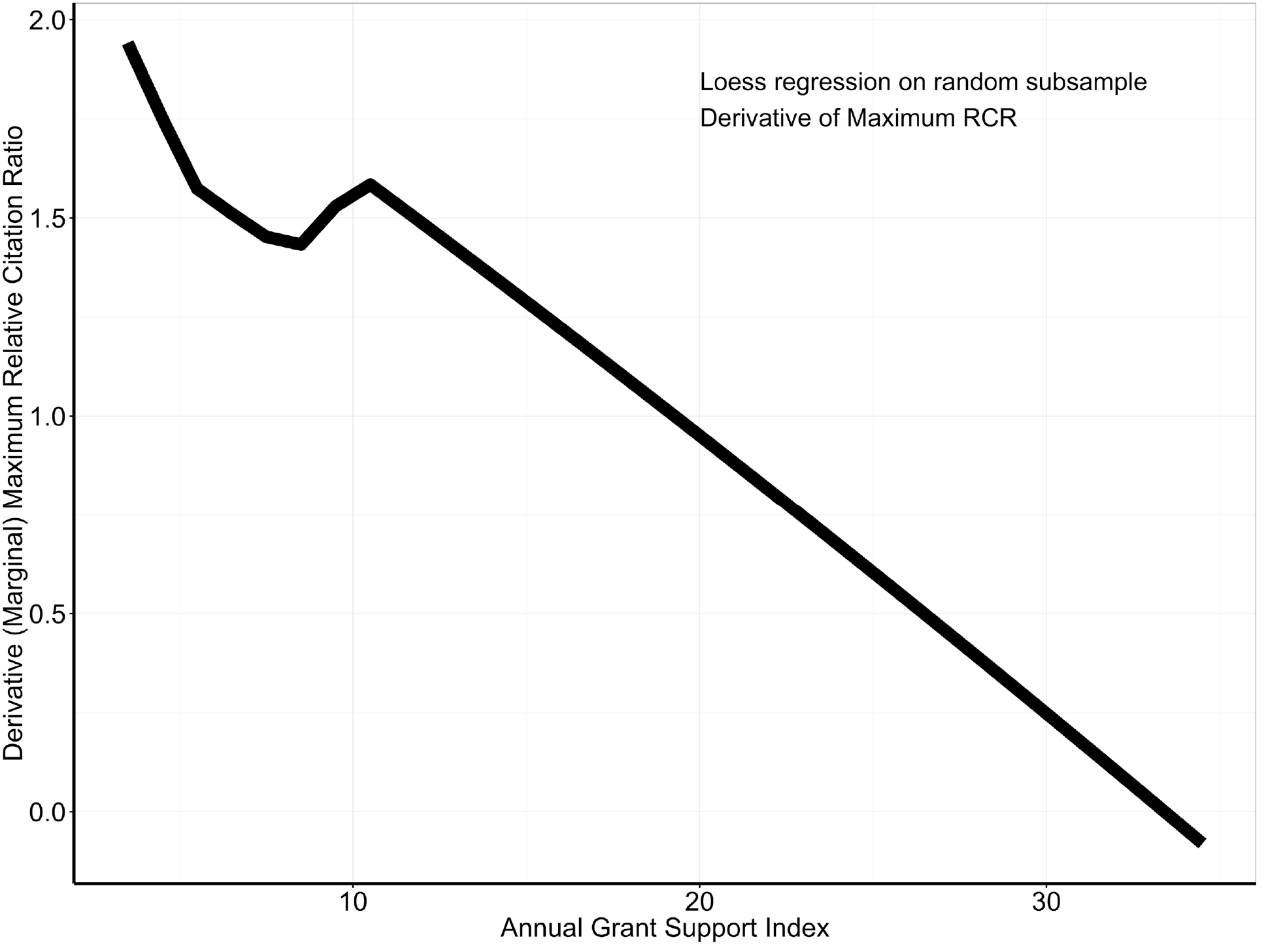
Marginal returns of maximum RCR by level of Grant Support – loess regression of random subsample. Association of the marginal returns of maximum Relative Citation Ratio (RCR) among all papers linked to a scientists’ grants with annual Grant Support Index (GSI). The curve shows the slope of the loess curve shown in Figure 1B and corresponds to the first derivative *dQ/dK* of the where *Q* is output as assessed by annual weighted RCR and *K* is input, here maximum RCR. Here *Q* and *K* are not log-transformed. Marginal returns fall as annual Grant Support Index increases.

Figure 3 and 4 are corresponding plots of the association of median RCR with annual GSI. As seen in Figure 4, marginal returns fall as annual GSI increases; negative returns are seen at annual GSI levels exceeding 21. Between annual GSI levels of 10 and 21, a proportional increase of input is associated with a lesser increase in output; beyond an annual GSI of 21, absolute output falls as annual GSI increases

**Figure 3:**
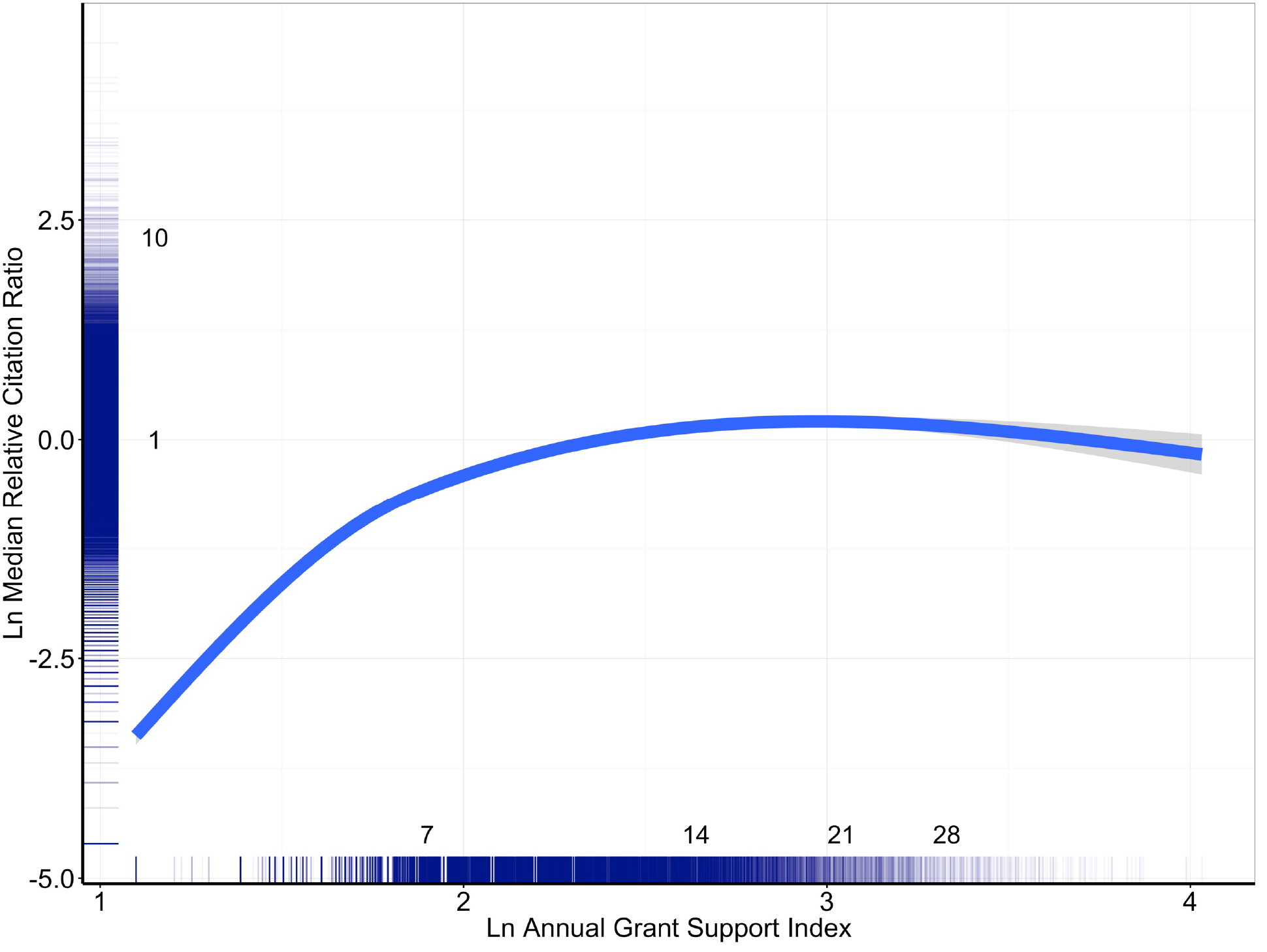
Median Relative Citation Ratio by Level of Grant Support. Association, as assessed by spline regression, of the median Relative Citation Ratio (RCR) among all papers linked to a scientists’ grants with annual Grant Support Index (GSI). Due to skewed distributions, both RCR and GSI values are natural log-transformed; the numbers inside the axes represent the raw, non-transformed values. An annual GSI value of 7 corresponds to ~ 1 R01 grant, while annual GSI values of 14 and 21 correspond to 2 and 3 R01 grants. The rugplots by the axes illustrate the log-normal distributions. Note the decreasing slope of the regression curve as annual GSI increases.

**Figure 4:**
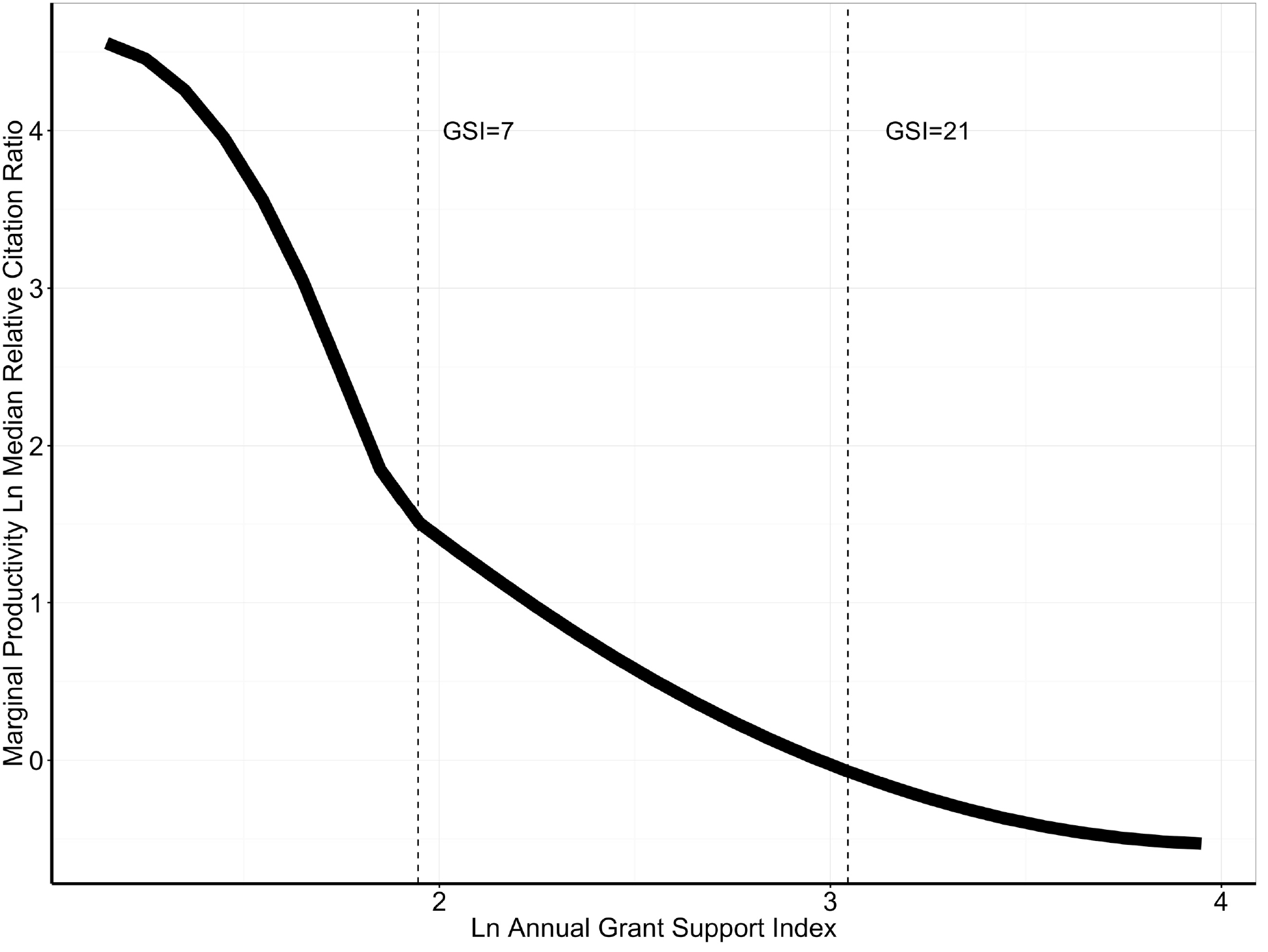
Marginal returns of median RCR by level of Grant Support. Association of the marginal returns of median Relative Citation Ratio (RCR) among all papers linked to a scientists’ grants with annual Grant Support Index (GSI). The curve shows the slope of the curve shown in Figure 3 and corresponds to the first derivative *d(ln(Q))/d(ln(K))* of the Cobb-Douglas function *ln(Q) ~ ln(α)* + *β(ln(K)),* where *Q* is output as assessed by median RCR and p is a coefficient linking the natural logarithm of input (K or *annual GSI)* with the natural logarithm of output, here median RCR. For convenience, vertical dashed lines are shown corresponding to GSI values of 7 (equivalent to 1 single-PI R01) and 21 (equivalent to 3 single–PI R01s). Marginal productivity falls as annual Grant Support Index increases. Values below zero, seen above annual GSI levels of 21, indicate negative returns.

Figures 5A, 5B, and 6A and 6B are corresponding spline and loess regression plots of the association of annual weighted RCR with annual GSI (along with first derivatives). Marginal returns fall with increasing annual GSI.

### Consideration of “elite” scientists

To assess these association among “elite” scientists, particularly those successful in securing additional funding, we stratified our sample by receipt of Howard Hughes Medical Institute (HHMI) funding between 1996 and 2014. Table 3 shows characteristics and outcomes as listed in Table 2; as might be expected, HHMI scientists received more NIH support for a longer time. They also published more papers that had higher RCR values. Figure 7 shows the associations of annual GSI with maximum (panel A), median (panel B), and annual weighted (panel C) RCR; decreasing marginal returns with higher annual GSI are evident for both HHMI and non-HHMI scientists.

**Table 3:**
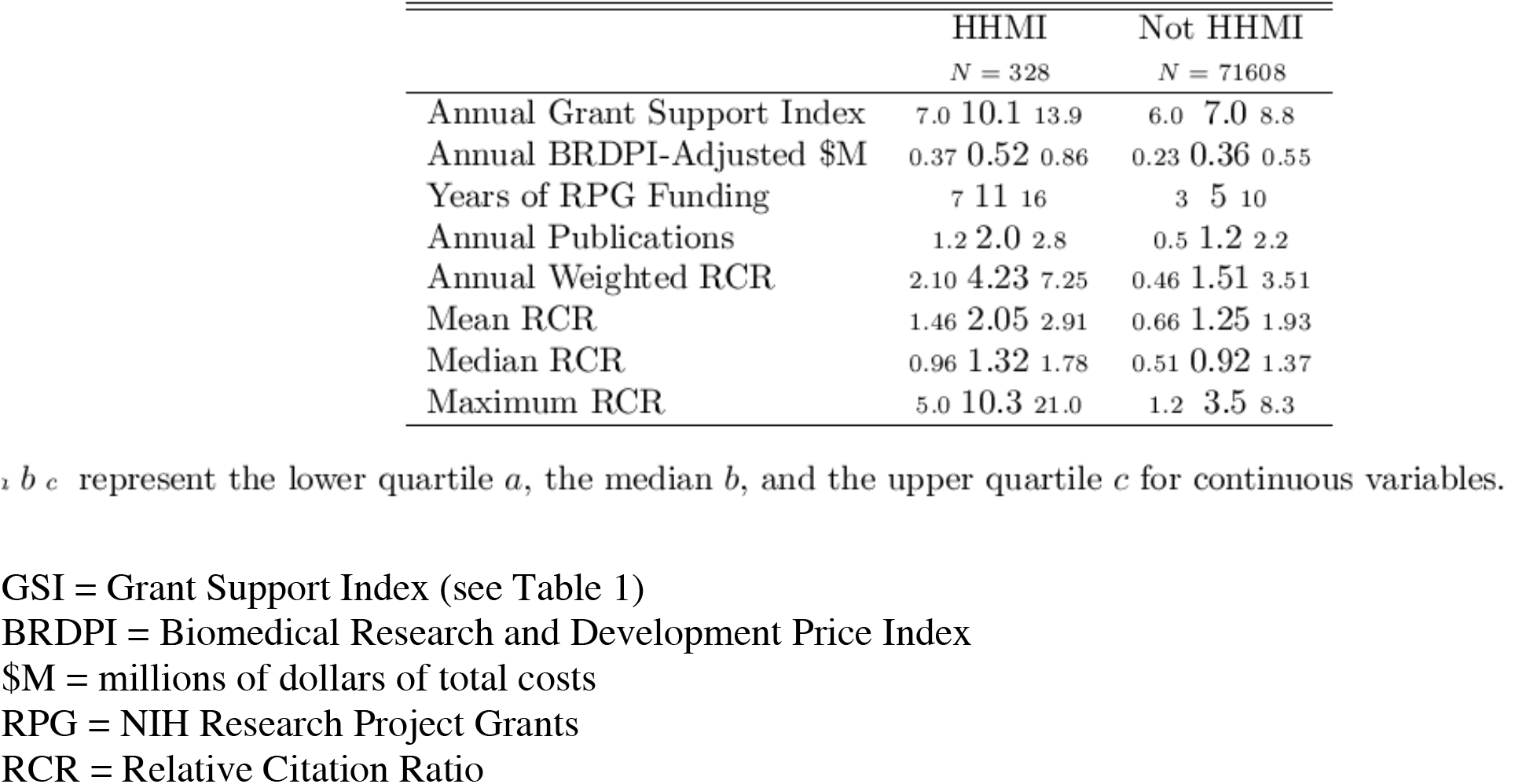
Characteristics and outputs of 71,936 NIH-supported scientists according to whether they received Howard Hughes Medical Institute (HHMI) funding between 1996 and 2014.

**Figure 5A:**
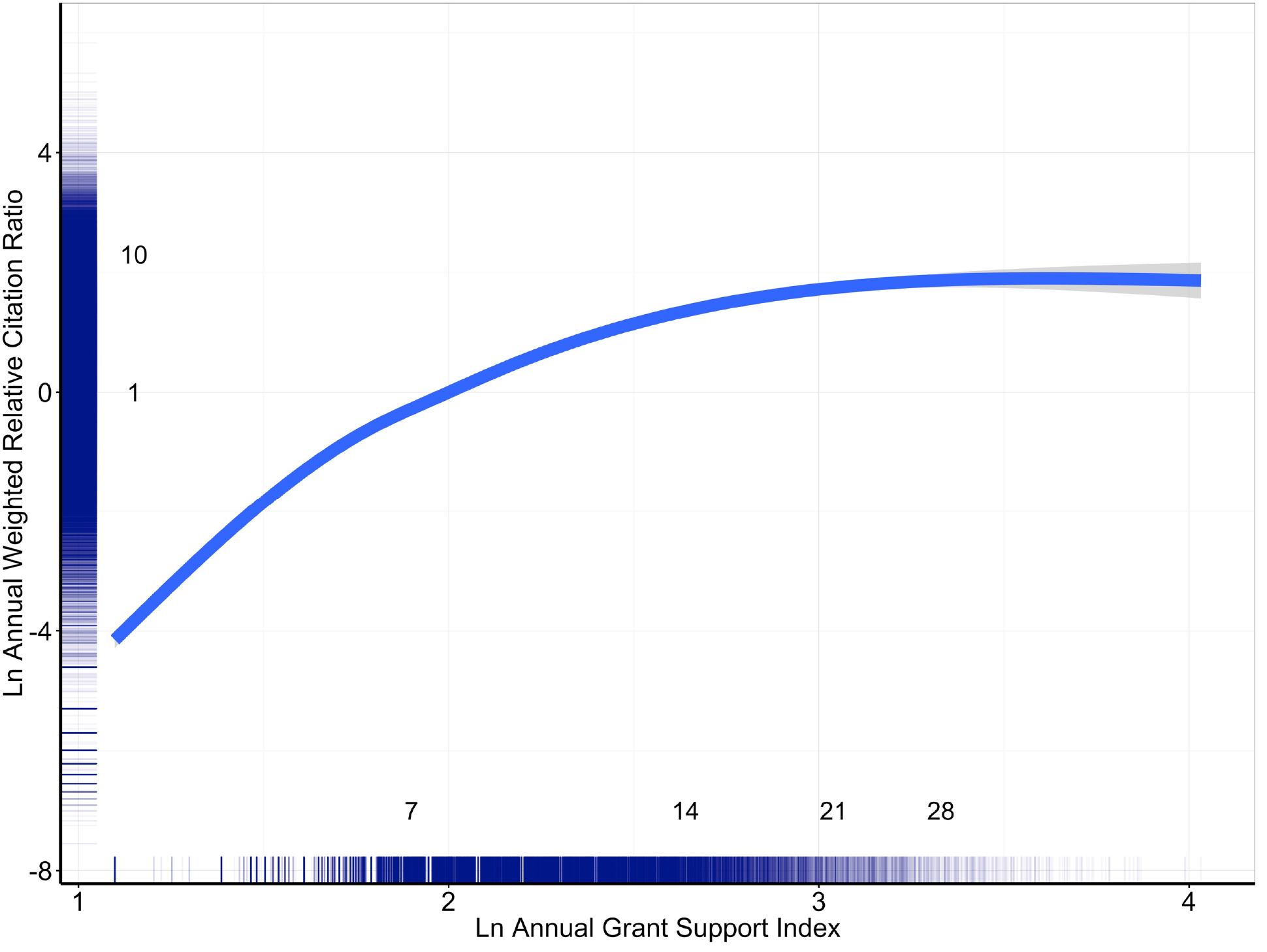
Annual Weighted Relative Citation Ratio by Level of Grant Support. Association, as assessed by spline regression, of the annual weighted Relative Citation Ratio (RCR) among all papers linked to a scientists’ grants with annual Grant Support Index (GSI). Due to skewed distributions, both RCR and GSI values are natural log-transformed; the numbers inside the axes represent the raw, non-transformed values. An annual GSI value of 7 corresponds to ~ 1 R01 grant, while annual GSI values of 14 and 21 correspond to 2 and 3 R01 grants. The rug-plots by the axes illustrate the log-normal distributions. Note the decreasing slope of the regression curve as annual GSI increases.

**Figure 5B:**
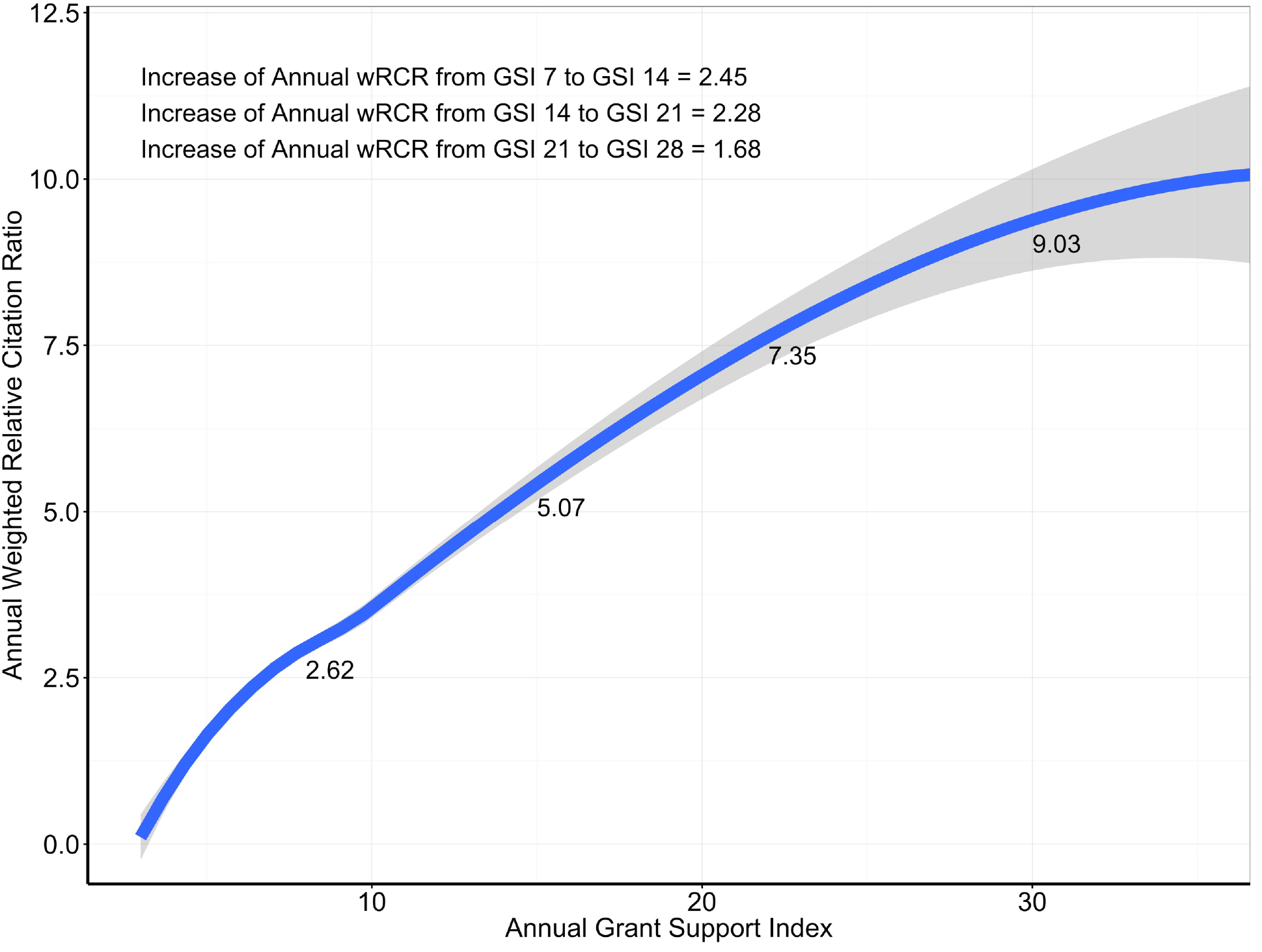
Annual Weighted Relative Citation Ratios and Grant Support – loess regression of random subsample. Association, as assessed by loess regression in a random subsample of 20,000, of the annual weighted Relative Citation Ratio (wRCR) among all papers linked to a scientists’ grants with annual Grant Support Index (GSI). Here values are not log-transformed. Loess fit values and changes are shown along the curve and in the upper left-hand corner to illustrate decreasing incremental returns with increasing levels of grant support. The decreasing instantaneous slope (by first derivative) is shown in Figure 6B.

**Figure 6A:**
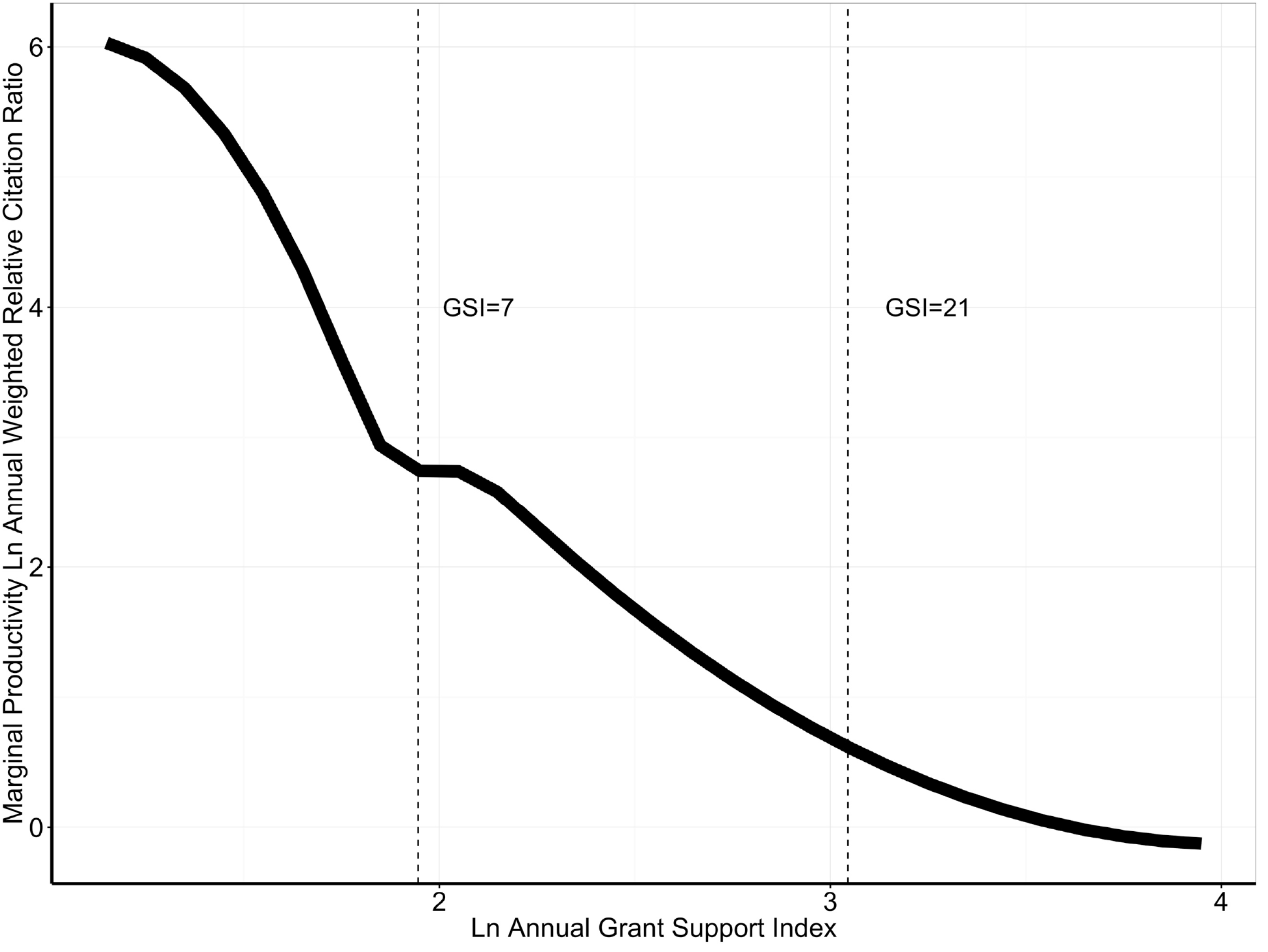
Marginal returns of annual weighted RCR by level of Grant Support. Association of the marginal returns of annual weighted Relative Citation Ratio (RCR) among all papers linked to a scientists’ grants with annual Grant Support Index (GSI). The curve shows the slope of the curve shown in Figure 5A and corresponds to the first derivative *d(ln(Q))/d(ln(K))* of the Cobb-Douglas function *ln(Q) ~ ln(α)* + *β(ln(k)),* where *Q* is output as assessed by annual weighted RCR and p is a coefficient linking the natural logarithm of input (K or *annual GSI)* with the natural logarithm of output, here annual weighted RCR. For convenience, vertical dashed lines are shown corresponding to GSI values of 7 (equivalent to 1 single-PI R01) and 21 (equivalent to 3 single-PI R01s). Marginal productivity falls as annual Grant Support Index increases.

**Figure 6B:**
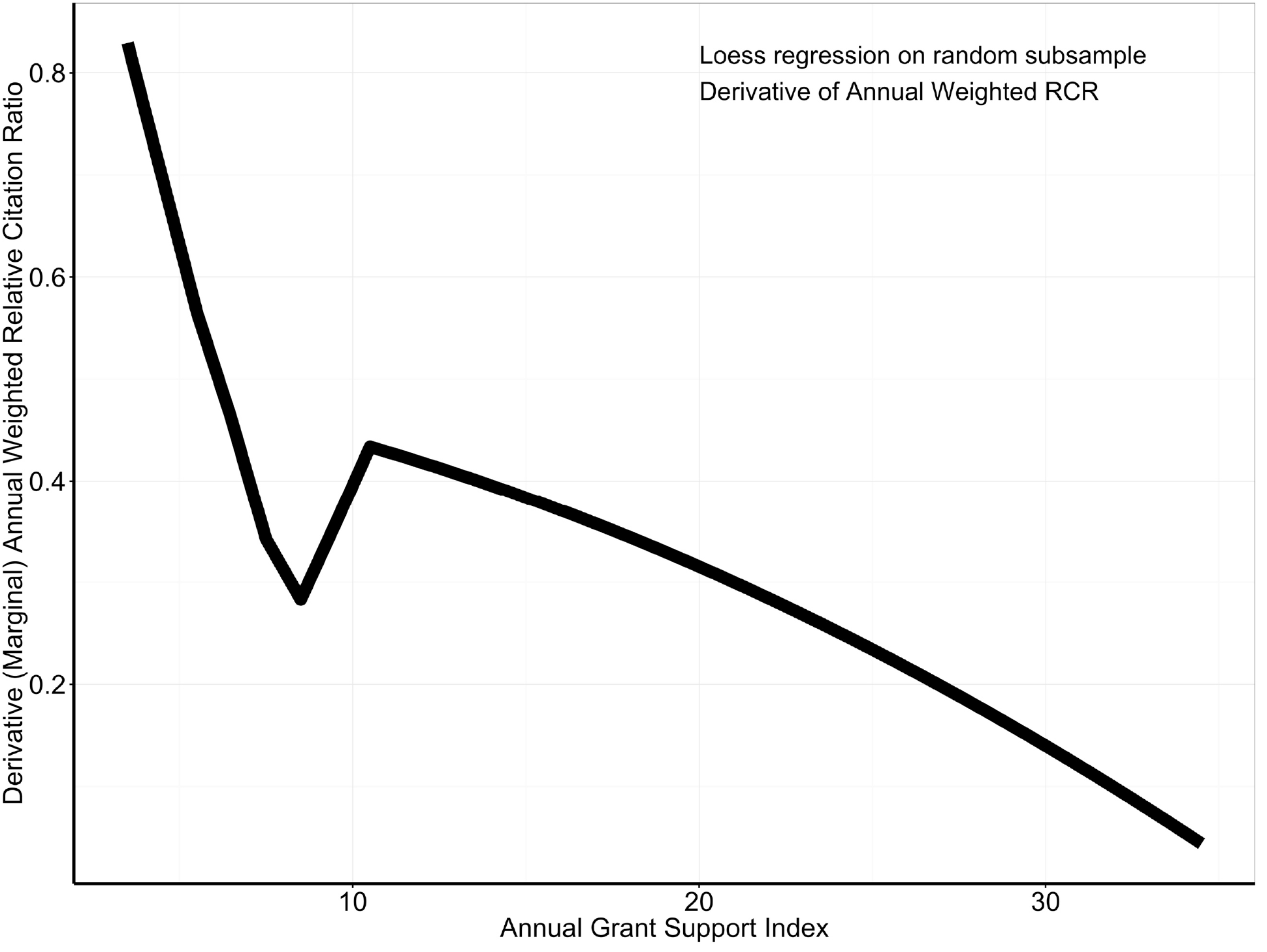
Marginal returns of annual weighted RCR by level of Grant Support – loess regression of random subsample. Association of the marginal returns of annual weighted Relative Citation Ratio (RCR) among all papers linked to a scientists’ grants with annual Grant Support Index (GSI). The curve shows the slope of the loess curve shown in Figure 5B and corresponds to the first derivative *dQ/dK* of the where *Q* is output as assessed by annual weighted RCR and *K* is input, here annual weighted RCR. Here *Q* and *K* are not log-transformed. Marginal returns fall as annual Grant Support Index increases.

**Figure 7:**
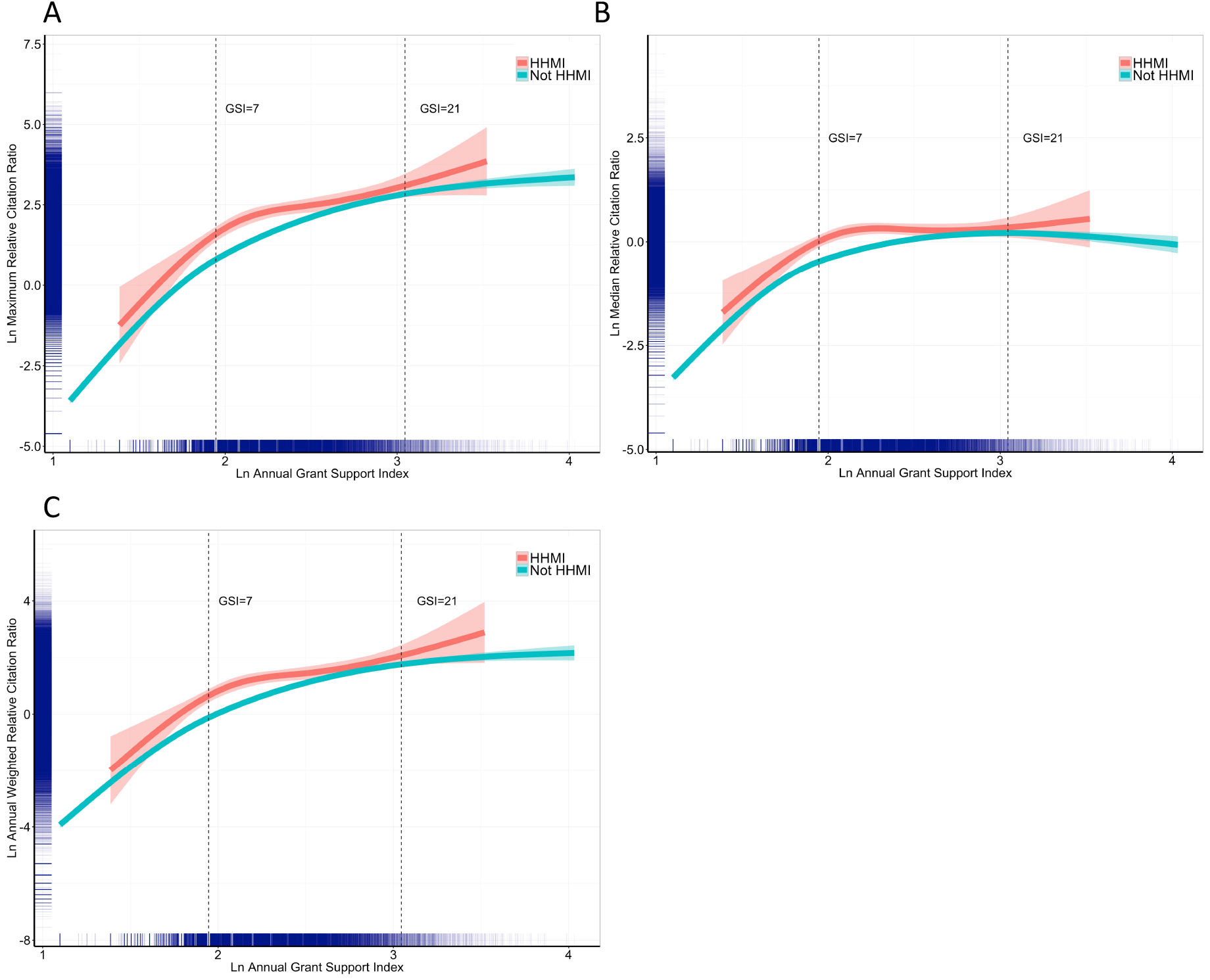
Associations of maximum (A), median (B), and annual weighted (C) Relative Citation Ratio according to annual GSI and HHMI funding status.

As another assessment of the impact of elite scientists, we identified the “top 50” who received at least 4 years of NIH funding by the 3 different RCR measures. Among the top 50 by maximum RCR, only 3 (6%) were scientists with annual GSI levels exceeding 21. Among the top 50 by median RCR, none had annual GSI levels exceeding 21. Among the top 50 by annual weighted RCR, only 7 (10%) had annual GSI levels exceeding 21.

### Secondary Analyses

We performed several other secondary analyses, including focusing on human–and non-human grants as well as grants from specific NIH Institutes (among them the National Cancer Institute, the National Institute of Allergy and Infectious Disease, the National Institute of General Medical Sciences, and the National Institute of Neurological Diseases and Stroke). In all cases, we saw similar patterns of decreasing marginal returns with increasing annual GSI.

Because of the complexity of multi-component awards and cooperative agreements, and because of misclassification errors in assigning scientists to sub-projects of those awards, we performed a secondary analysis focusing on scientists’ support from single-component R awards only (in other words, excluding P and U awards and excluding all subprojects, N=65,574 scientists). Figure 8 shows similar patterns of falling marginal returns for maximum (panel A), median (panel B), and annual weighted (panel C) RCR as annual GSI increases.

**Figure 8:**
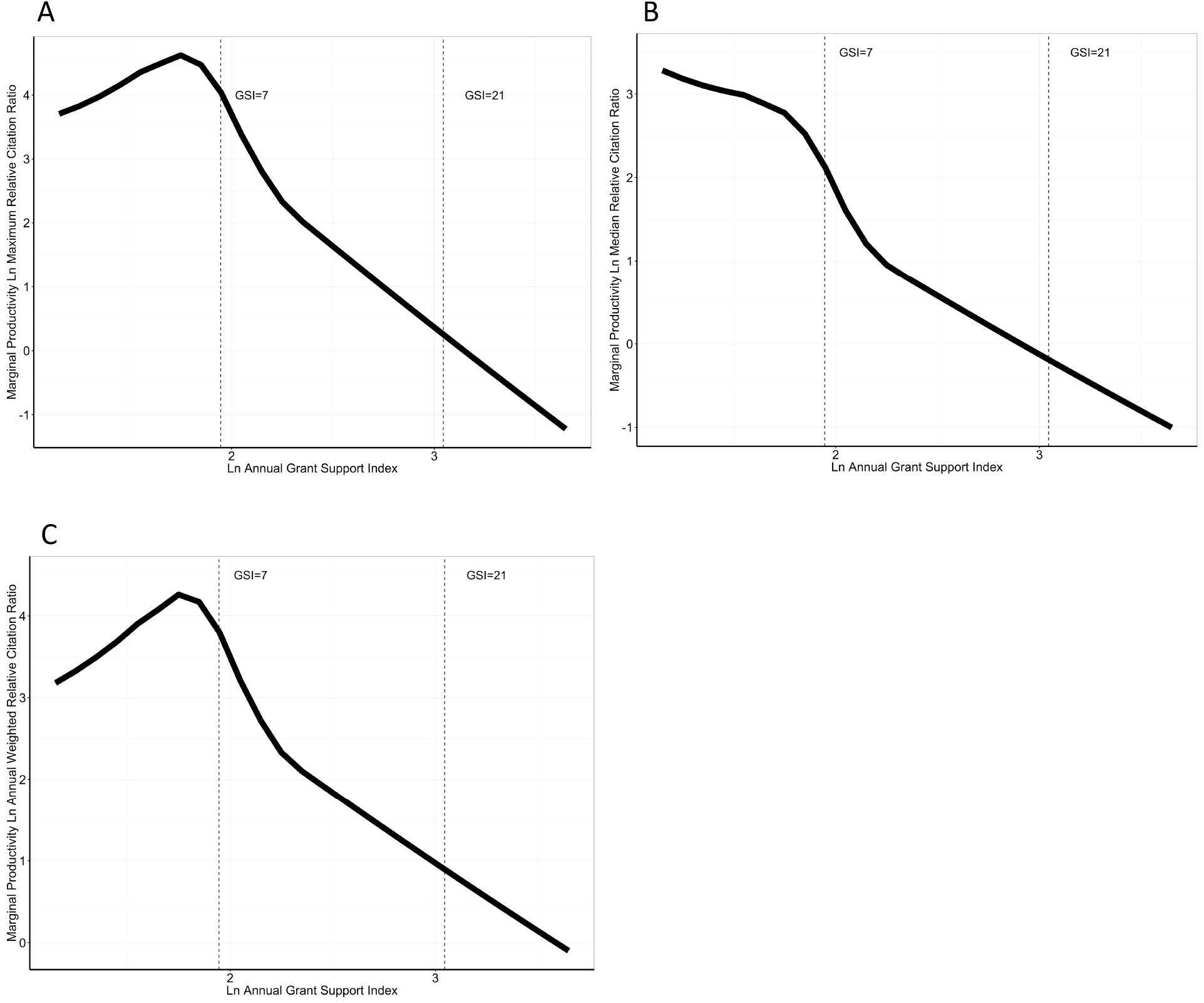
Secondary analyses limited to grant support on single-component “R” grants. Association of the marginal returns of maximum (panel A), median (panel B), and annual weighted (panel C) Relative Citation Ratio (RCR) among all papers linked to a scientists’ grants with annual Grant Support Index (GSI). These analyses were based on R grants only. The curves shows the slope of the curve corresponding to the first derivative of the Cobb-Douglas function *ln(Q) ~ ln(α)* + *β(ln(annual GSI)),* where Q is output as assessed by maximum, median, or annual weighted RCR and b is a coefficient linking the natural logarithm of input (annual GSI) with the natural logarithm of output. For convenience, vertical dashed lines are shown corresponding to GSI values of 7 (equivalent to 1 single-PI R01) and 21 (equivalent to 3 single–PI R01s).

Finally, we looked at marginal returns as a function of dollars of funding (as done in previously published reports) among all scientists and among all RPG grants. Figure 9 shows rising and falling marginal returns for maximum (panel A), median (panel B), and annual weighted (panel C), with peaks at ~$400,000 per year.

**Figure 9:**
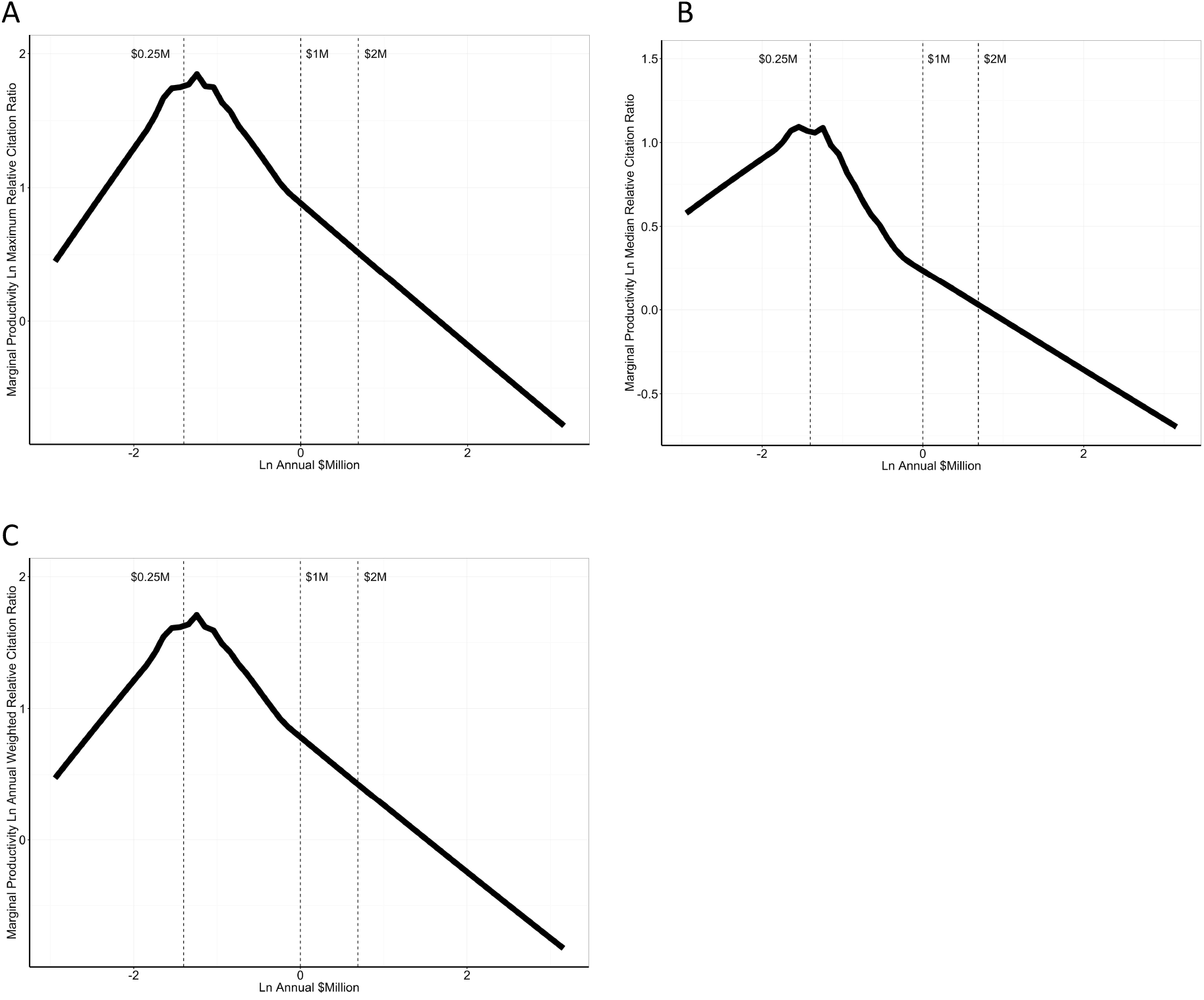
Marginal returns by levels of financial support. Association of the marginal returns of maximum (panel A), median (panel B), and annual weighted (panel C) Relative Citation Ratio (RCR) among all papers linked to a scientists’ grants with annual funding in BRDPI-adjusted dollars. The curves shows the slope of the curve corresponding to the first derivative of the Cobb-Douglas function *ln(Q) ~ ln(α)* + *β(ln(annual funding)),* where Q is output as assessed by maximum, median, or annual weighted RCR and b is a coefficient linking the natural logarithm of input (annual funding) with the natural logarithm of output. For convenience, vertical dashed lines are shown corresponding to funding values of $250,000, $1 million, and $2 million.

## Discussion

Despite using a different approach to measuring levels of laboratory support, we confirm the findings of others who have claimed that research output does not necessarily scale to input. Previous reports have primarily focused on financial or personnel measures^2–8^, and have described decreasing marginal returns – that is output that does not scale proportionally to input. One group used a regression discontinuity design, and consistent with our results, found that receipt of an R01 grant (usually on top of existing funding) led to a modest 7% increase in productivity.^13^ In that study, the increment in productivity was greater in younger scientists (age < 45 years), perhaps because they had lesser baseline funding.

Because different fields of science entail different cost structures, we developed a “Research Commitment Index” or “Grant Support Index” to describe research support that is relatively independent of field or inherent costing challenges. Even with this measure, we find that by 3 measures – Relative Citation Ratio, median Relative Ratio, and annual weighted Relative Citation Ratio^9^ – marginal returns decrease with higher levels of grant support. Thus, we confirm with data Alberts’ et al contention that “at some point returns per dollar diminish.”^1^

There are important limitations in our analyses. We could not measure additional non–NIH resources, such as “hard money” that might come with endowed chairs or university–supported tenured faculty positions. Nonetheless, we found similar patterns of decreasing marginal returns among HHMI–supported faculty. We could not be particularly granular across fields of study, though the Relative Citation Ratio seeks to account for differences across disciplines. We did not have data on the make–up of scientific teams supporting each scientists; more recent work from our group^14^ suggests that prospective assessments of team size and make–up may be feasible. Nonetheless, prior work has found evidence of decreasing marginal returns with larger team size.^2^ NIH grant mechanisms and funding structures are complex, and, as an example, we couldn’t tease the effects of infrastructure and training support. We only focused on citation metrics, and will leave to future work to assess the associations of NIH grant support on training, development of technologies, translation, and sharing of resources, these and other measures have been advocated as legitimate measures of research success.^15^ The GSI point system was developed by NIH staff using a semi–quantitative approach, and of note was written down before any of these outcomes analyses were performed. We were aware of questions about how best to classify work on parent and subprojects for multicomponent grants; nonetheless, secondary analyses focused on singlecomponent R grants yielded similar findings (Figure 8). Since <u>we publicized a proposed policy</u> on grant limits to enable funding more new investigators, we have heard much feedback and are considering several changes which may affect the findings presented here. We present the findings here based on the older point scale since that point scale was agreed upon before undertaking any outcomes analyses.

Despite these limitations, we found that we could reproduce others’ findings of decreasing marginal returns even by using a measure of research support distinct from money or personnel. Some have argued that the phenomena of decreasing marginal returns offers funders an opportunity to enhance the likely productivity of their research portfolio by <u>seeking to fund as many scientists as possible.</u>^16^ Furthermore, as others have argued^3^, science is often unpredictable^11,17^, and therefore funders increase the likelihood of supporting transformative discoveries by seeking ways to fund more scientists across the spectrum of their careers.^11^ Funders’ ability to support as many meritorious scientists as possible is arguably more important than ever given rapidly improving technologies and worsening hypercompetition.^18^

## Supplementary materials

All data and annotated R statistical code used to generate the Tables and Figures in this report are available at Bioarxiv.org.

### Supplemental Figures

**Figure S1:**
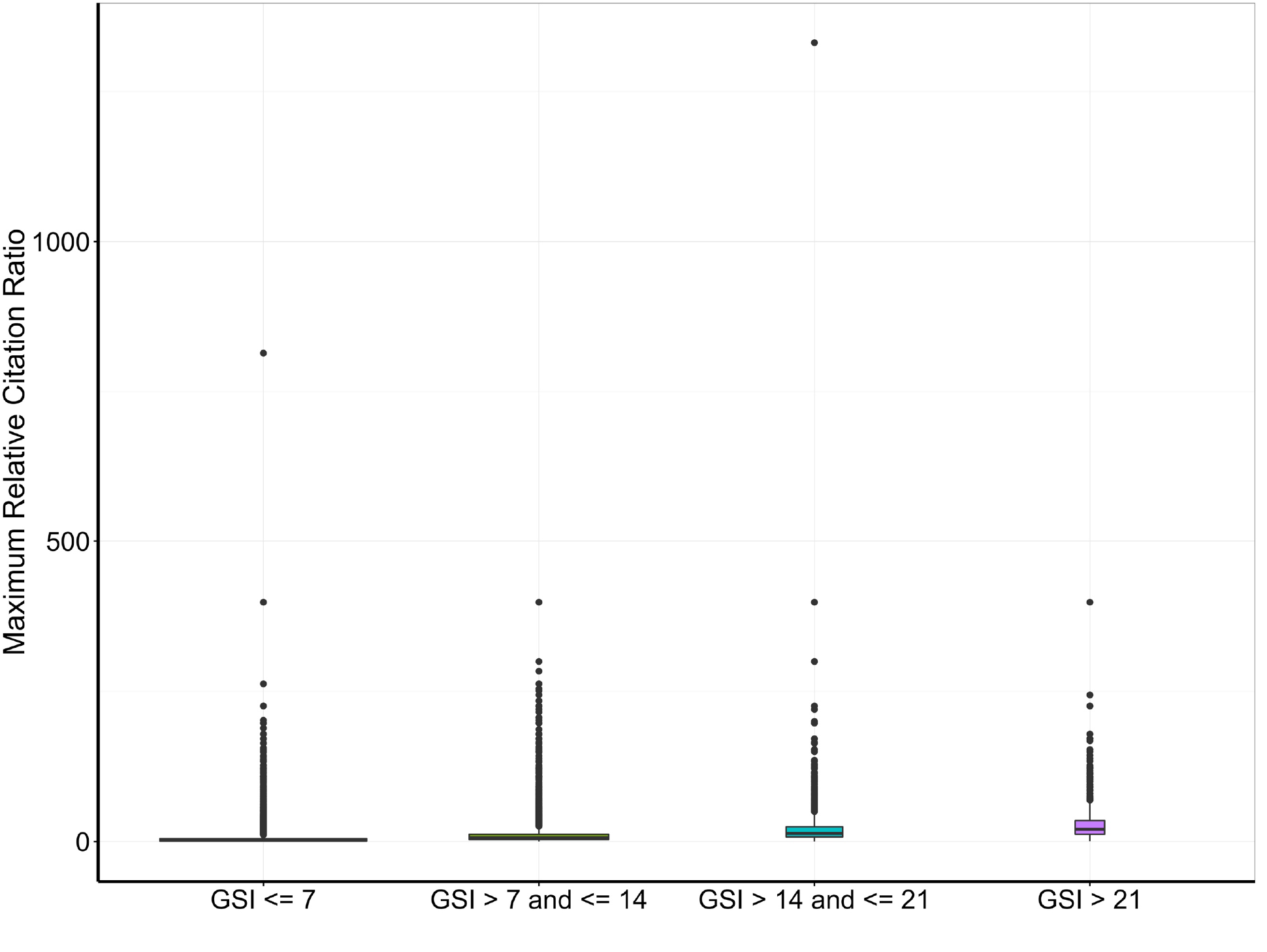
Boxplots of maximum RCR according to the annual GSI categories shown in Table 2.

**Figure S2:**
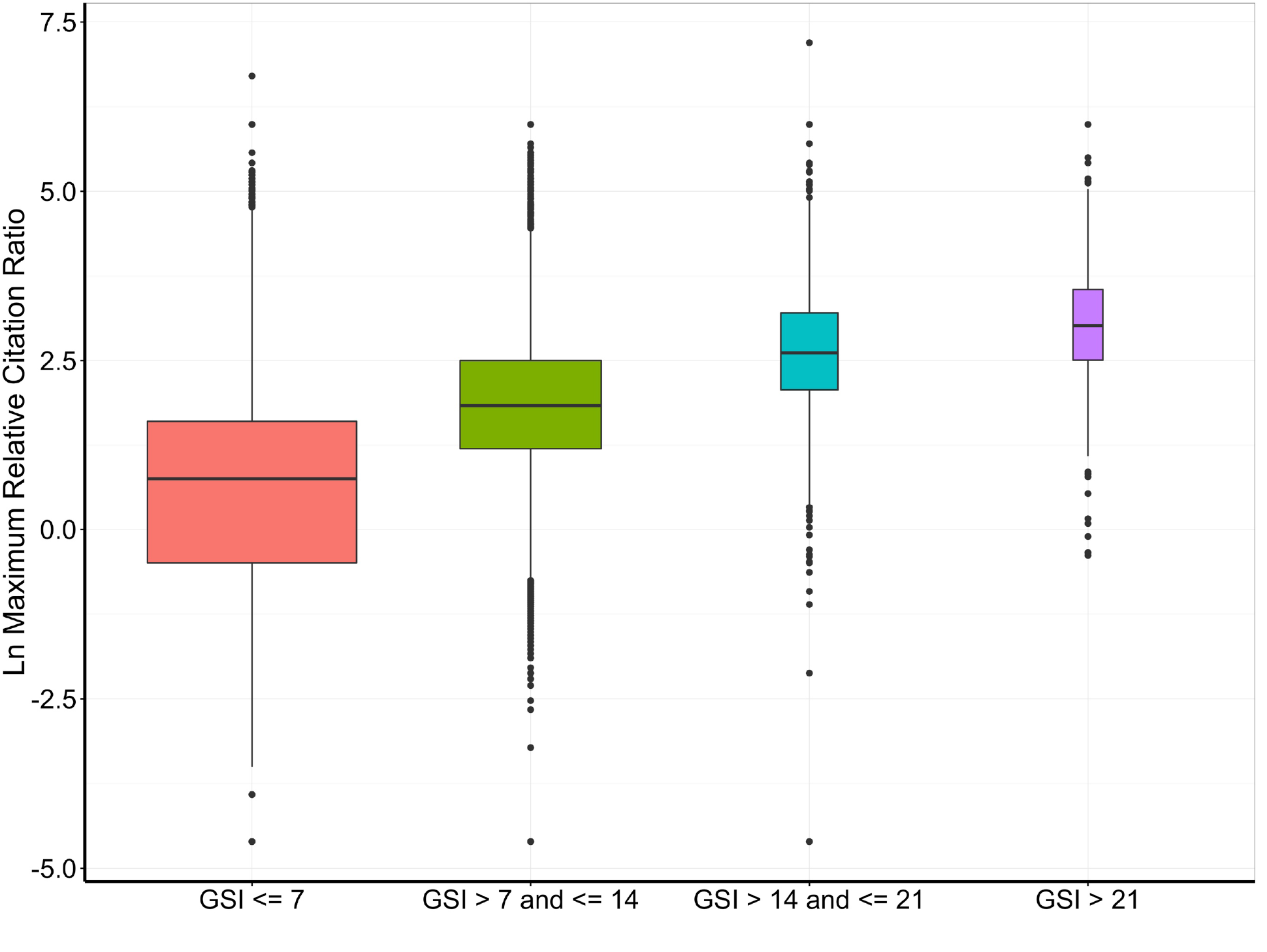
Same data as in Figure S1, but natural log-transformed

**Figure S3:**
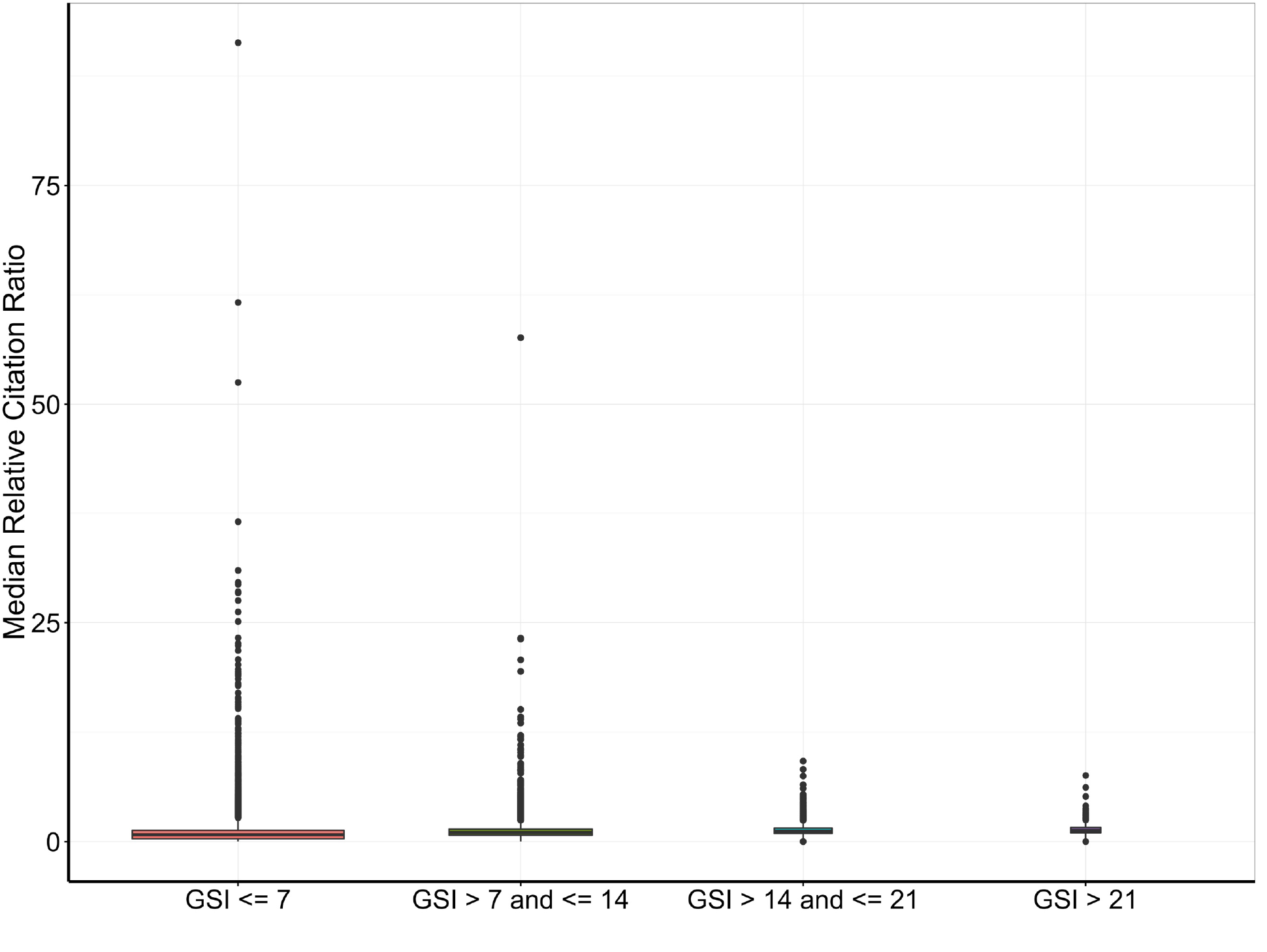
Boxplots of median RCR according to the annual GSI categories shown in Table 2.

**Figure S4:**
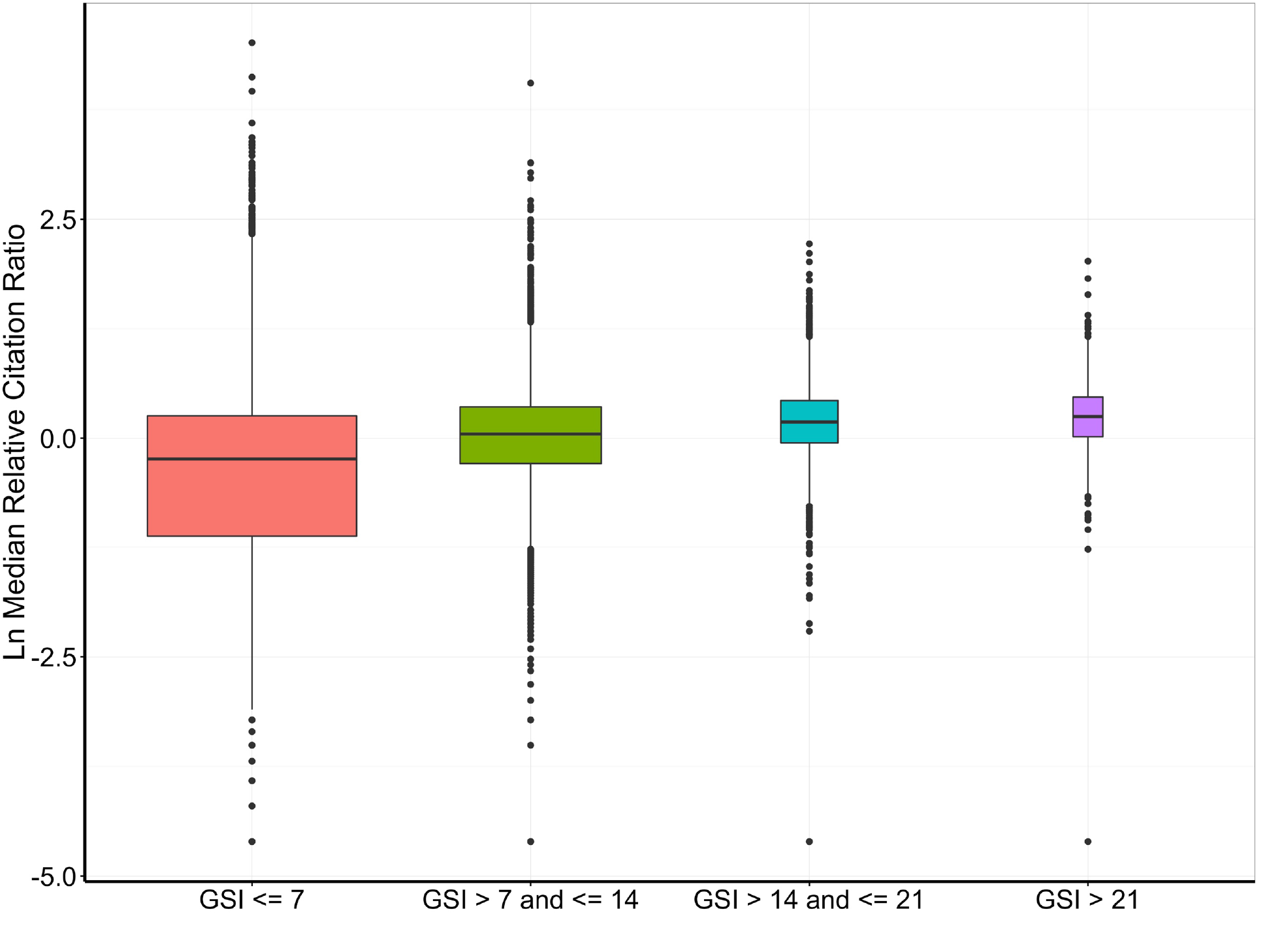
Same data as in Figure S3, but natural-log transformed.

**Figure S5:**
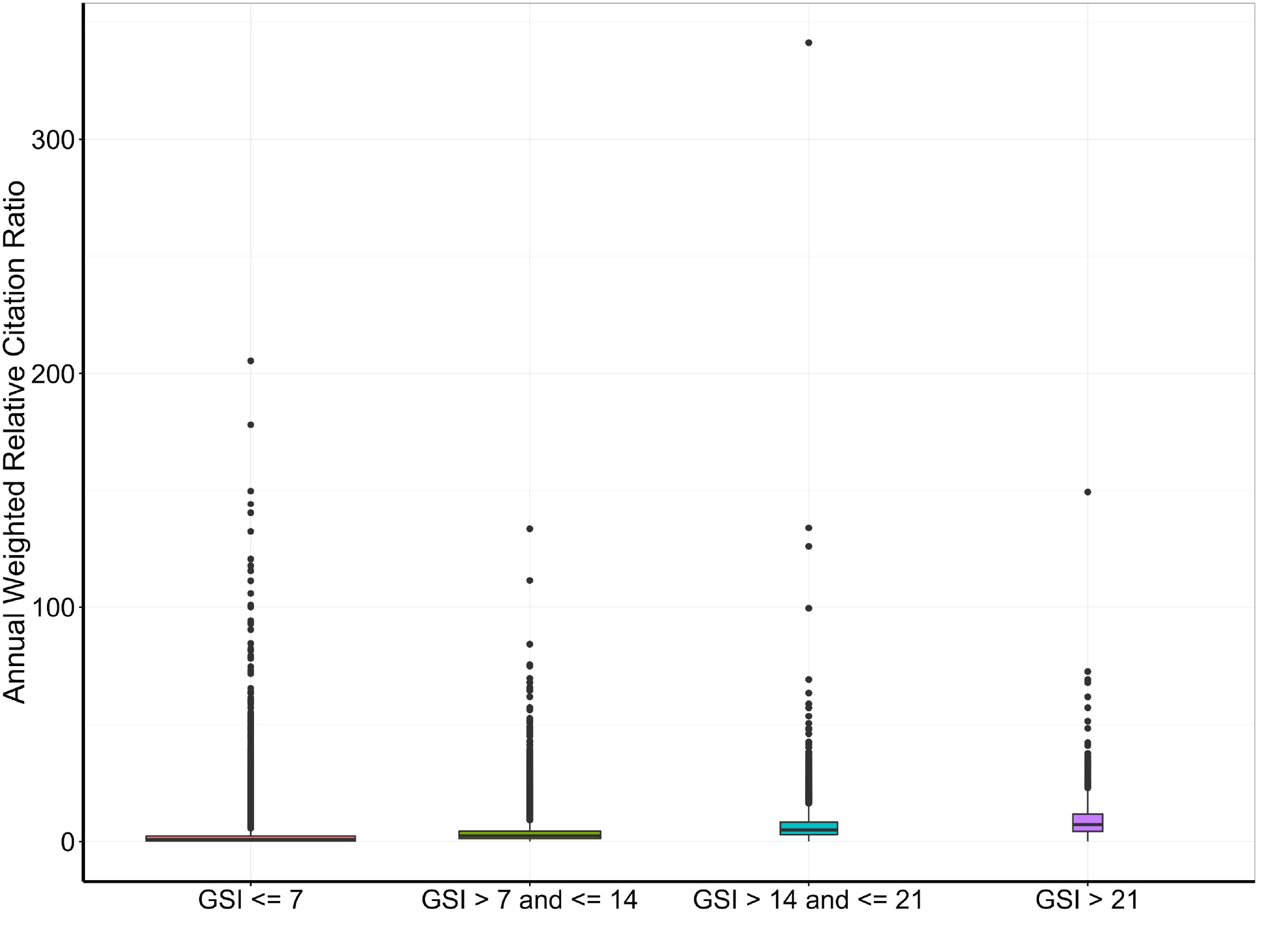
Boxplots of annual weighted RCR according to the annual GSI categories shown in Table 2.

**Figure S6:**
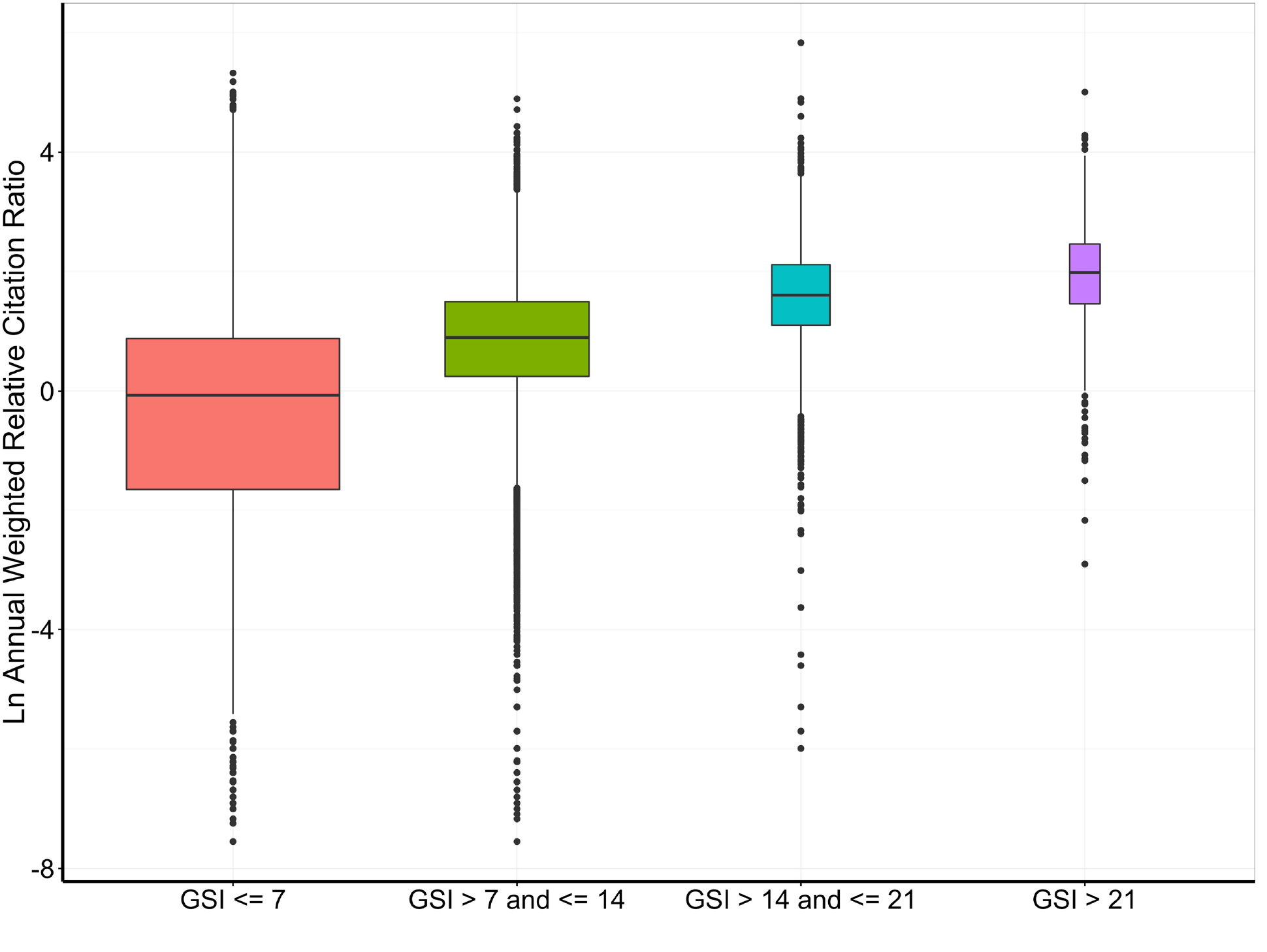
Same data as in Figure S5, but natural log-transformed

